# Intestinal infection results in impaired lung innate immunity to secondary respiratory infection

**DOI:** 10.1101/2020.08.03.235457

**Authors:** Shubhanshi Trivedi, Allie H. Grossmann, Owen Jensen, Mark J. Cody, Taylor A. Wahlig, Paula Hayakawa Serpa, Charles Langelier, Kristi J. Warren, Christian C. Yost, Daniel T. Leung

## Abstract

**Background:** Pneumonia and diarrhea are among the leading causes of death worldwide, and epidemiological studies have demonstrated that diarrhea is associated with an increased risk of subsequent pneumonia. Our aim was to determine the impact of intestinal infection on innate immune responses in the lung.

**Methods:** Using a mouse model of intestinal infection by *Salmonella enterica* serovar Typhimurium (*S*. Typhimurium (*ST*)), we investigated how infection in the gut compartment can modulate immunity in the lungs and impact susceptibility to bacterial (*Klebsiella pneumoniae* (*KP*)) challenge.

**Results:** We found alterations in frequencies of innate immune cells in lungs of intestinally-infected mice compared to uninfected mice. On subsequent challenge with *K. pneumoniae* we found that mice with prior intestinal infection have higher lung bacterial burden and inflammation, increased neutrophil margination, and neutrophil extracellular traps (NETs), but lower overall numbers of neutrophils, compared to mice without prior intestinal infection. Total numbers of dendritic cells, innate-like T cells, and natural killer cells were not different between mice with and without prior intestinal infection.

**Conclusions:** Together, these results suggest that intestinal infection impacts lung innate immune responses, most notably neutrophil characteristics, potentially resulting in increased susceptibility to secondary pneumonia.

**Article summary:** We show, in a mouse model, that prior intestinal infection with *Salmonella Typhimurium* leads to increased susceptibility to respiratory *Klebsiella pneumoniae infection*, which is associated with altered neutrophil responses.

## Background

Diarrhea and pneumonia are among the leading causes of death worldwide. In children alone, these diseases combine to kill ~1.4 million each year, with the majority of these deaths occurring in lower and middle-income countries [1]. Epidemiological studies have shown that children are at an increased risk of pneumonia following a diarrheal episode [2, 3]. However, the immunological mechanisms behind an increased susceptibility to such secondary respiratory infections are not well understood.

Although the gastrointestinal and respiratory tracts have different environments and functions, there is emerging data showing cross-talk between these two mucosal sites in chronic inflammatory diseases such as inflammatory bowel disease (IBD) and asthma [4]. Additionally, there is emerging evidence that the intestinal microbiota plays a role in host defense against bacterial pneumonia [5]. The gastrointestinal and respiratory tracts share the same embryonic origin and have common components of the mucosal immune system such as an epithelial barrier, submucosal lymphoid tissue, the production of IgA and defensins, and the presence of innate lymphocytes and dendritic cells [6]. Notably, several innate-like leukocytes, such as Mucosal Associated Invariant T (MAIT) cells, invariant Natural Killer T (iNKT) cells, gamma delta T (γδ T) cells, dendritic cells (DC) and neutrophils, have the capacity to circulate between tissues, and play important roles in both respiratory and intestinal tract immunity [7, 8].

How intestinal infection impacts immunity in the lung is not known. In this study, we examine the impact of intestinal infection on the immune response in the lungs of mice using an established model of intestinal infections. We found that mice infected with *S*. Typhimurium (*ST*) have increased susceptibility to respiratory *Klebsiella pneumoniae* (*KP*) infection compared to mice without prior intestinal infection. Prior intestinal infection modulated effector cells of innate immunity in the lung, contributing to respiratory immune dysregulation and a higher *KP* bacterial burden.

## Methods

### Mice and inoculations

Six to eight week old female C57BL/6J wild type mice were obtained from Jackson Laboratories. All animals were maintained and experiments were performed in accordance with University of Utah and Institutional Animal Care and Use Committee approved guidelines (protocol 17-01011). The animals were kept at a constant temperature (25°C) with unlimited access to pellet diet and water in a room with a 12 h light/dark cycle. All animals were monitored daily and infected animals were scored for the signs of clinical illness severity [9]. Animals were ethically euthanized using CO_2._

For the experiments, all mice were pretreated with streptomycin as described previously [10]. Briefly, water and food were withdrawn 4 hours before oral gavage treatment with 7.5 mg of streptomycin in 100 µl HBSS. Afterward, animals were supplied with water and food ad libitum. At 20 h after streptomycin treatment, water and food were withdrawn again for 4 hours before mice were gavaged with 10^4^ CFU of *ST* (100 μl suspension in PBS) or treated with sterile PBS (control). Thereafter, drinking water and food ad libitum was offered immediately. Six days post infection, mice were euthanized and lungs removed for analysis.

For *KP* challenge experiments, six days post *ST* infection, isoflurane anesthetized mice were inoculated intranasally with high inoculum ~ 10^10^ CFU *KP* in a 50 µl volume.Inoculated mice were euthanized at 18 h for bacterial load enumeration and immune assessment (details of bacterial strains and dosing justifications in Supplementary Methods).

### Lung Histology

After sacrifice, mouse lungs were infused with 10% neutral buffered formalin via the trachea, fixed in formalin overnight, dehydrated in 70% ethyl alcohol and embedded in paraffin. 4 µm sections were stained with haematoxylin and eosin and analysed by a board certified anatomic pathologist (A.H.G.). Samples were blinded prior to histopathologic analysis.

### Lung Neutrophil Extracellular Trap Assessment

Paraffin imbedded mouse lung were cut to 8 μm thickness on a microtome. Sections were deparaffinized and rehydrated using xylene and decreasing ethanol concentration washes. Heat induced epitope antigen retrieval of lung sections were processed in a 2100 Retriever Thermal Processor (Electron Microscopy Sciences) containing Citrate buffer pH 6.0 solution. Sections were incubated for 10 minutes with 0.1% Triton-X-100 and blocked with 10% Donkey Serum for 1 hour at RT. Antibodies for citrullinated Histone H3 (Abcam) and myeloperoxidase (MPO; R&D Systems) were incubated at 1:100 dilution in 10% donkey serum overnight at 4°C. After washing sections with PBS, secondary rabbit and goat antibodies conjugated to Alexa Fluor 488 or Alexa Fluor 546, respectively, along with DAPI nuclear stain were incubated on sections for 90 minutes at 4°C. Sections were washed and coverslips were adhered with aqueous mounting medium (Dako, S3025). Images were acquired on an Olympus FV3000 Confocal Laser Scanning Microscope. FluoView software (Olympus) and ImageJ Fiji (NIH) were used for image processing and analysis. We quantified NET formation on the images using a standardized grid system [11]. Briefly, we placed a standardized grid on randomly selected high-power field images (n=5 field images/sample). The number of times that any NET crossed a grid line was tallied.

### Lung mononuclear cell isolation

For lung digestion and preparation of single cell suspensions, lungs were perfused using 5 mL PBS, aseptically harvested from euthanized mice and kept in RPMI with 10% Fetal Bovine Serum (FBS). Lungs were dissociated using the mouse Lung Dissociation Kit (Miltenyi Biotec) and the gentleMACS Dissociator (Miltenyi Biotec). Cells were then passed through a 70 µm cell strainer and washed. Red blood cells were lysed with red blood cell lysis buffer. Lung mononuclear cells were then washed twice.

### Bacterial Load quantification

*KP* bacterial load was determined by plating ten-fold serial dilutions of the lung homogenates onto MacConkey agar plates (Sigma-Aldrich). The plates were incubated at 37°C overnight before bacterial CFUs were determined by colony counts.

### Mouse inflammation quantification

Lungs homogenates were filtered on 70 μm cell strainers and centrifuged at 300 × g for 5 min. Supernatants were stored at −80°C for cytokine content analysis. Lung cytokine levels were assessed from the supernatant samples via LEGENDplex kit (mouse inflammation panel 13-plex; BioLegend) per manufacturer’s instructions. Cytokine levels were acquired using a FACSCanto II flow cytometer (BD Biosciences), and analyses were performed using LEGENDplex data analysis software (BioLegend).

### Tetramer and antibody surface-staining of lung single cell suspensions

From each animal, 1-2 million cell aliquots were prepared and stained with the fixable viability dye eFluor™ 780 (eBioscience) for 15 min at room temperature (RT) to exclude dead cells, and washed with PBS + 2% FBS and incubated with anti-mouse CD16/CD32 Fc Block antibody (BD), for 20 min at 4 °C. Cells were then stained for 30 min with fluorescence-conjugated antibodies at either RT or 4 °C (detailed in Supplementary Methods). A total of 10^6^ gated events per sample were collected using the BD Fortessa flow cytometer, and results analyzed using FlowJo 10.4.2 software.

### Statistical analysis

GraphPad Prism 8 software was used for statistical analysis. The Mann-Whitney *U* test was used for comparison between uninfected and *ST* infected groups and between mice with and without prior intestinal infection. Results were presented as mean ± standard deviation, and *p* < 0.05 was considered statistically significant.

## Results

### 1. Mice with prior *ST* intestinal infection have increased lung bacterial load, sickness score, and susceptibility to respiratory *KP* infection

To test whether prior intestinal infection increases the susceptibility to respiratory *KP* infection, we evaluated survival, body weight loss, sickness score, and bacterial burden in lungs post *KP* challenge (Fig 1A). Compared to *ST* infected mice, which had a survival rate of 90-100%, and *K. pneumonia* infected mice, which had a survival of 60-80%, mice with prior *ST* intestinal infection had a survival rate of 0-30% at 120 hours post *KP* infection (Fig 1B). Interestingly, although there were no statistically significant differences in body weight loss between mice with or without *ST* infection with *KP* challenge (Fig S1), mice with prior intestinal infection had a higher sickness score (Fig 1C). This sickness scoring system included hunched posture, ruffled fur, decreased movement and altered respiratory rates and quality of breaths (Fig 1C). Of note, mice with only *ST* intestinal infection showed no weight loss and had reduced sickness scores. When bacterial burden was evaluated at 18 and 33 hours post *KP* infection, we found significantly higher lung bacterial burden in mice with prior intestinal infection compared to mice without prior intestinal infection (Fig 1D). Mice with prior intestinal infection also showed significantly higher lung bacterial burden at 33 hours compared to 18 hours (Fig 1D). We could not isolate any *ST* from lung homogenates of mice with intestinal infection. In addition, in a subset of animals, we performed metagenomic next generation sequencing (Supplementary Methods) and observed the presence of *ST* in stool samples and *KP* in bronchoalveolar lavage (BAL) fluid (supplementary table 1).

**Figure 1:**
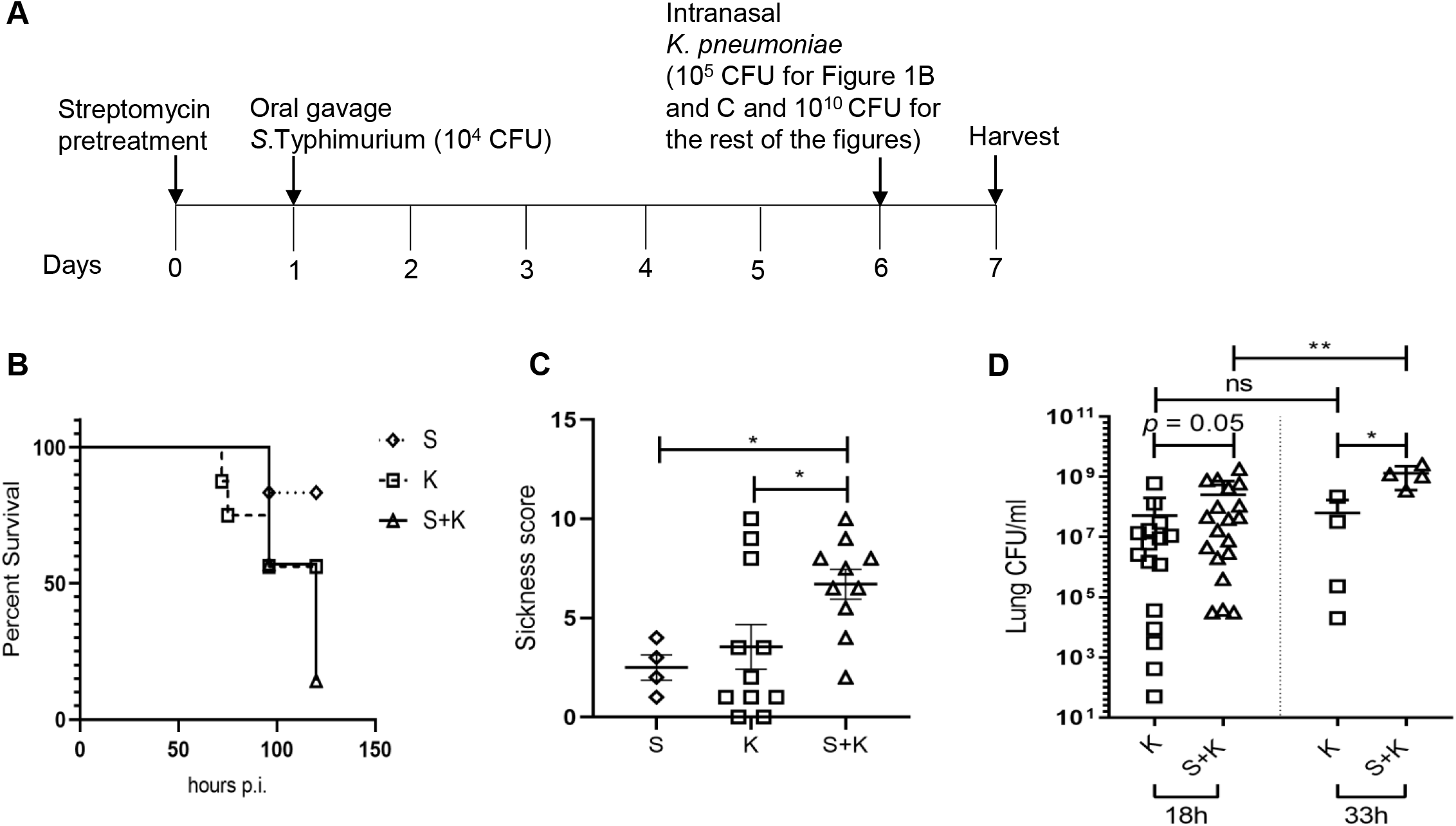
Mice with prior *ST* intestinal infection have increased susceptibility to respiratory *KP* infection. (A) Timeline of this study. (B) Kaplan-Meier survival curves of mice infected with *ST* intestinal infection (S), *KP infection* only (K) and mice with prior intestinal *ST* infection and challenged with *KP* infection (S+K). Statistical analysis was performed using log-rank (Mantel-Cox) test, p = 0.07 and log-rank test for significant trend, p = 0.02 (C) Sickness score plots where data represent cumulative results of two independent experiments (n = 4-6 for S, n = 11 for K and n = 10 for S+K) and mean ± SD, * denotes p < 0.05. Statistical analysis was performed using Kruskal-Wallis test followed by Dunn’s multiple comparison test. (D) *KP* bacterial load determined in lungs at 18 hours post *KP* infection in both K and S+K groups. Data represents cumulative results of three independent experiments (n = 16 for K and n = 19 for S+K). Lung bacterial loads also determined at 33 hours in both groups (n = 4 for each group, N = 1 experiment). Data represent mean ± SD and statistical analysis was determined using Mann-Whitney test; * denotes p < 0.05, ** denotes p < 0.01.

### 2. Mice with prior *ST* intestinal infection have increased lung inflammation from subsequent respiratory *KP* infection

We next examined the degree of lung inflammation by histological analysis of tissue sections from all four groups of mice. Histopathological analysis revealed mixed interstitial inflammatory consolidations in mice with *KP* respiratory infection and increased microabcess formation with pyknotic neutrophils in mice co-infected with *ST* and *KP* (Fig 2A and supplementary table 2). Uninfected mice and mice with intestinal infection showed normal alveolar and interstitial lung histology (Fig 2A). Upon challenge with *KP* respiratory infection, mice with prior *ST* intestinal infection also showed marked intravascular clustering of polymorphonuclear neutrophils (PMNs), with increased margination, necrotic cluster formation and extravasation (Fig 2B and Fig S2). Intravascular neutrophil clustering was noticeably absent in the other treatment groups. Furthermore, lung sections from mice with intestinal infection, and mice with both intestinal and respiratory infection, showed scattered microthrombi in capillary-sized vessels whereas mice with *KP* respiratory infection (with no prior intestinal infection) showed no microthrombi formation (Fig S3).

**Figure 2:**
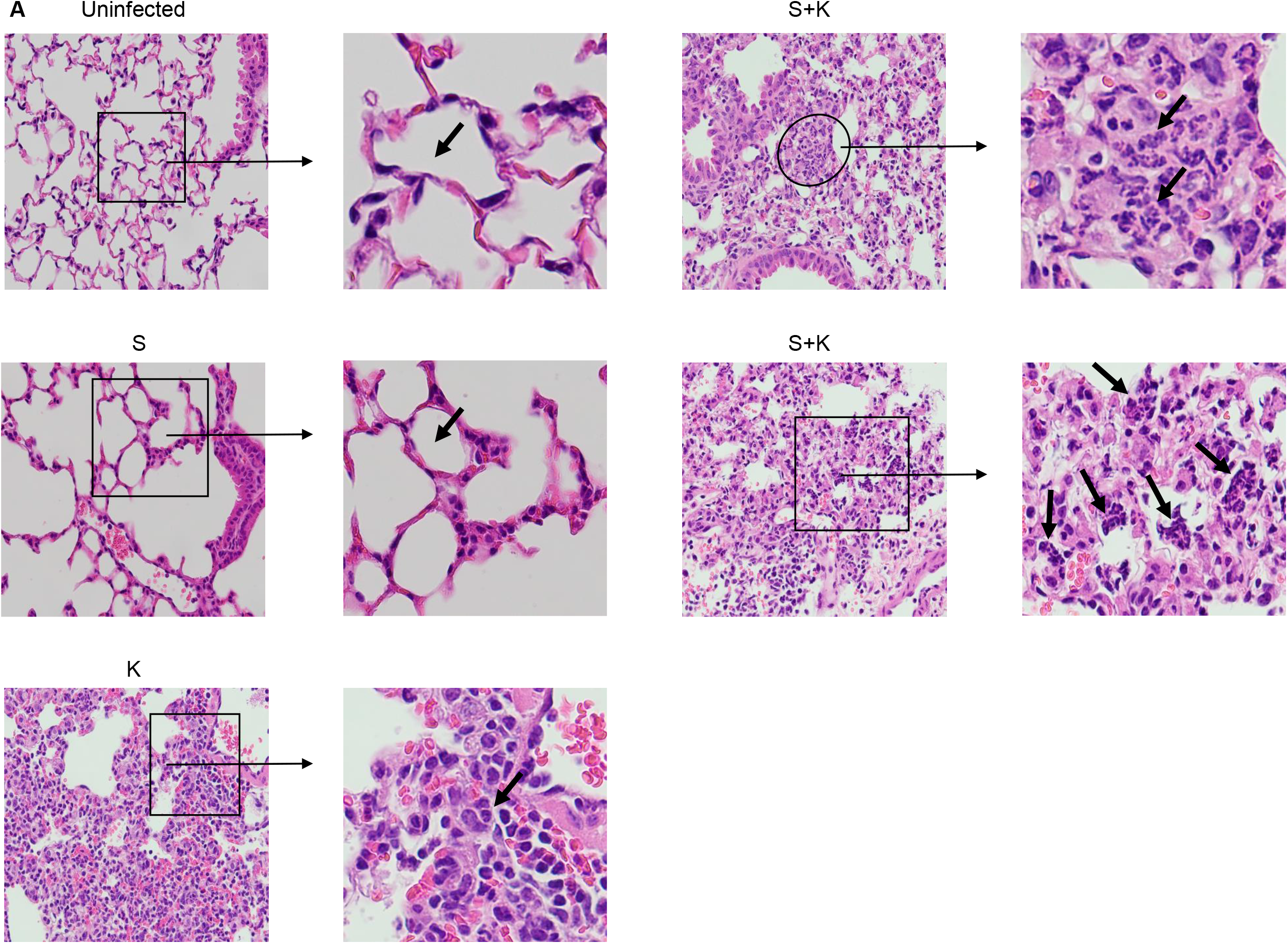
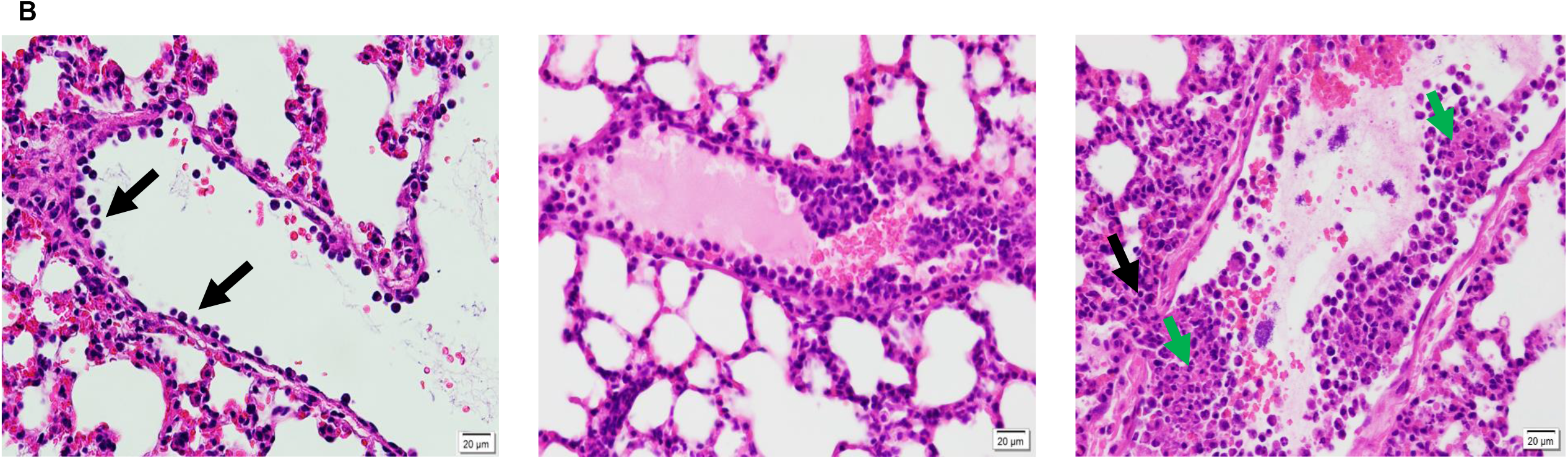
Mice with prior *ST* intestinal infection have increased lung inflammation from subsequent respiratory *KP* infection, characterized by microabcess, pyknotic neutrophil clusters and margination, compared to those with no prior intestinal infection. (A) Representative images (400x magnification, scale bar = 20 µm) of lung sections stained with hematoxylin/eosin, insets shows digital magnification of original image. In uninfected mice and in mice with intestinal infection only (S), black arrows show normal lung parenchyma. Mice with respiratory infection only (K) show mixed interstitial inflammatory consolidations, and black arrow shows a normal neutrophil. In mice with prior intestinal ST infection and challenged with KP infection (S+K), microabcess are circled and black arrows highlight clusters of pyknotic neutrophils. (B) Representative images (H & E, 400x magnification, scale bar = 20 µm) of lung sections of S + K infected mice. Lung vessels in S + K infected mice showing neutrophil margination (left, black arrows), margination and clustering (middle), necrotic clusters (right, green arrow) and extravasation (right, black arrow). Data represents cumulative results of two independent experiments (n = 2 for UI, n = 10 for S group, n = 9 for K group and n = 8 for S+K group).

### 3. Mice with prior *ST* intestinal infection have altered lung cytokine profiles after *KP* challenge

The delicate balance between pro- and anti-inflammatory cytokines is crucial in containing pathogens and maintaining tissue repair and homeostasis in the lung [11]. Prior studies have shown the importance of cytokines such as IFN-γ in recruitment of neutrophils to the lung tissue [12, 13]. We investigated whether intestinal infection affected cytokine production in lung homogenates, and whether prior intestinal infection affected this cytokine response to intranasal *KP* challenge. Our results indicate that compared with the uninfected control group, *ST* infected mice had significantly higher lung levels of IFN-γ, MCP-1 and IL-1β (Fig 3 A-C). While Thy1-expressing natural killer (NK) cells [14] and NKp46^+^ ILC3 [14] cells are commonly thought as the sources of IFN-γ, there is a mounting evidence that neutrophils are a prominent cellular source of IFN-γ during the innate phase of *ST*-induced colitis [15]. It is possible that large numbers of primed neutrophils traffick to lungs after intestinal infection, contribute to cytokine production and increase the potential for neutrophil-mediated pathology or NET formation upon secondary infection [8]. Furthermore, following *KP* challenge, mice with prior intestinal infection had higher levels of IFN-γ and lower levels of GM-CSF cytokine production in lung homogenates compared to those without prior intestinal infection (Fig 3A and D). Levels of IL-23, IL-1α, TNFα, IL-12p70, IL-10, IL-6, IL-27, IL-17A and IFN-β were not significantly different between mice with and without prior intestinal infection (Fig S4).

**Figure 3:**
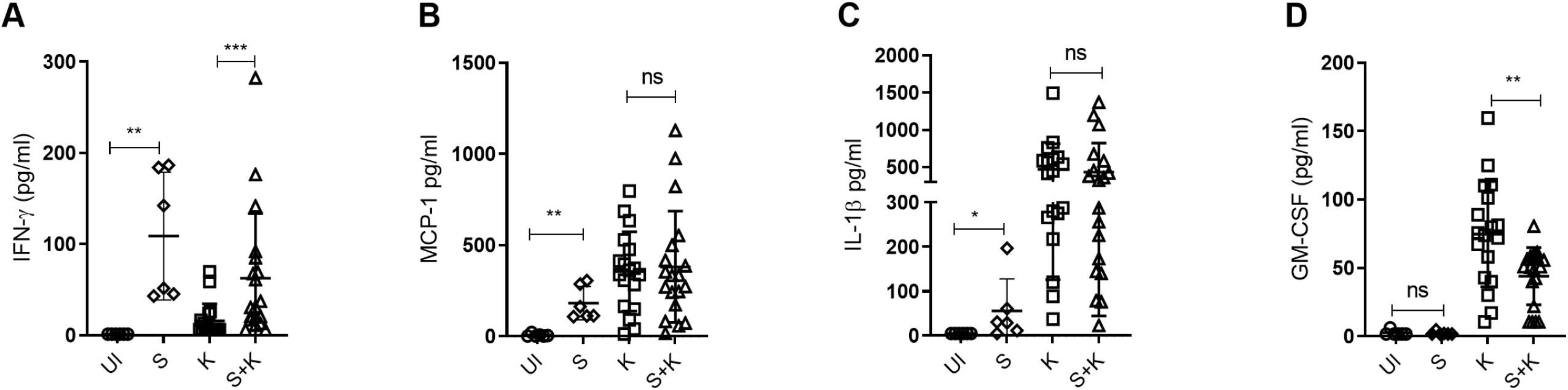
Mice with prior *ST* intestinal infection have altered lung cytokine profiles, and altered cytokine responses to respiratory *KP* infection, compared to those with no prior intestinal infection. Levels of lung inflammatory cytokines (A) IFN-γ (minimum detection limit (MDL) = 2.44 pg/ml)), (B) MCP-1 (MDL = 2.44 pg/ml), (C) IL-1β (MDL = 2.44 pg/ml) and (D) GM-CSF (MDL = 2.44 pg/ml) were assessed 18 hours post *KP* challenge via bead-based LEGENDplex mouse inflammation panel 13-plex assay. Data represents one experiment for uninfected mice (n = 6) and *ST* infected mice (n = 6) and cumulative results of three independent experiments for K (n = 18) and S+K group (n = 19). Error bars represents mean ± SD and significance was determined by Mann-Whitney tests.

### 4. Mice with prior *ST* intestinal infection have lower numbers of neutrophils in the lungs after *KP* challenge

We next examined the impact of intestinal infection on innate cellular responses in the lung, and also its effect on such responses to *KP* respiratory challenge. We analyzed changes in major innate lung leukocytes (plasmacytoid dendritic cells (pDCs), monocyte-derived dendritic cells (moDCs), CD103+ DCs, neutrophils, alveolar macrophages (AMs) and interstitial macrophages (IMs)) pre-and post-*KP* challenge in mice infected with *ST* (gating strategy in Fig S5A). Consistent with previous studies [9], we observed rapid and robust recruitment of neutrophils to the lungs at 18 hours following *KP* infection compared to uninfected controls. Interestingly, mice with prior intestinal *ST* infection had significantly lower frequencies and total number of lung neutrophils following *KP* challenge compared to mice infected with *KP* alone (Fig 4A). Furthermore, results indicated that frequencies of pDCs increased and moDCs decreased in the lungs of *ST* intestinally infected mice compared to uninfected controls (Fig 4B and C). Following intranasal *KP* challenge, we found a marked increase in frequencies of pDCs and significantly lower frequencies of moDCs in mice with prior *ST* intestinal infection compared to those without prior intestinal infection (Fig 4B and C). No significant differences were observed in frequencies of CD103^+^ DCs (Fig 4D) between *KP* infected mice with and without prior *ST* intestinal infection. Total numbers of pDCs, moDCs, CD103^+^ DCs, AMs or IMs were not different in mice with prior intestinal infection compared to those without prior *ST* intestinal infection (Fig 4 and Fig S6). We also found higher numbers of neutrophils, pDCs and IMs and lower numbers of CD103^+^ DCs in mice with only *ST* intestinal infection compared to uninfected mice (Fig 4 and Fig S6).

**Figure 4:**
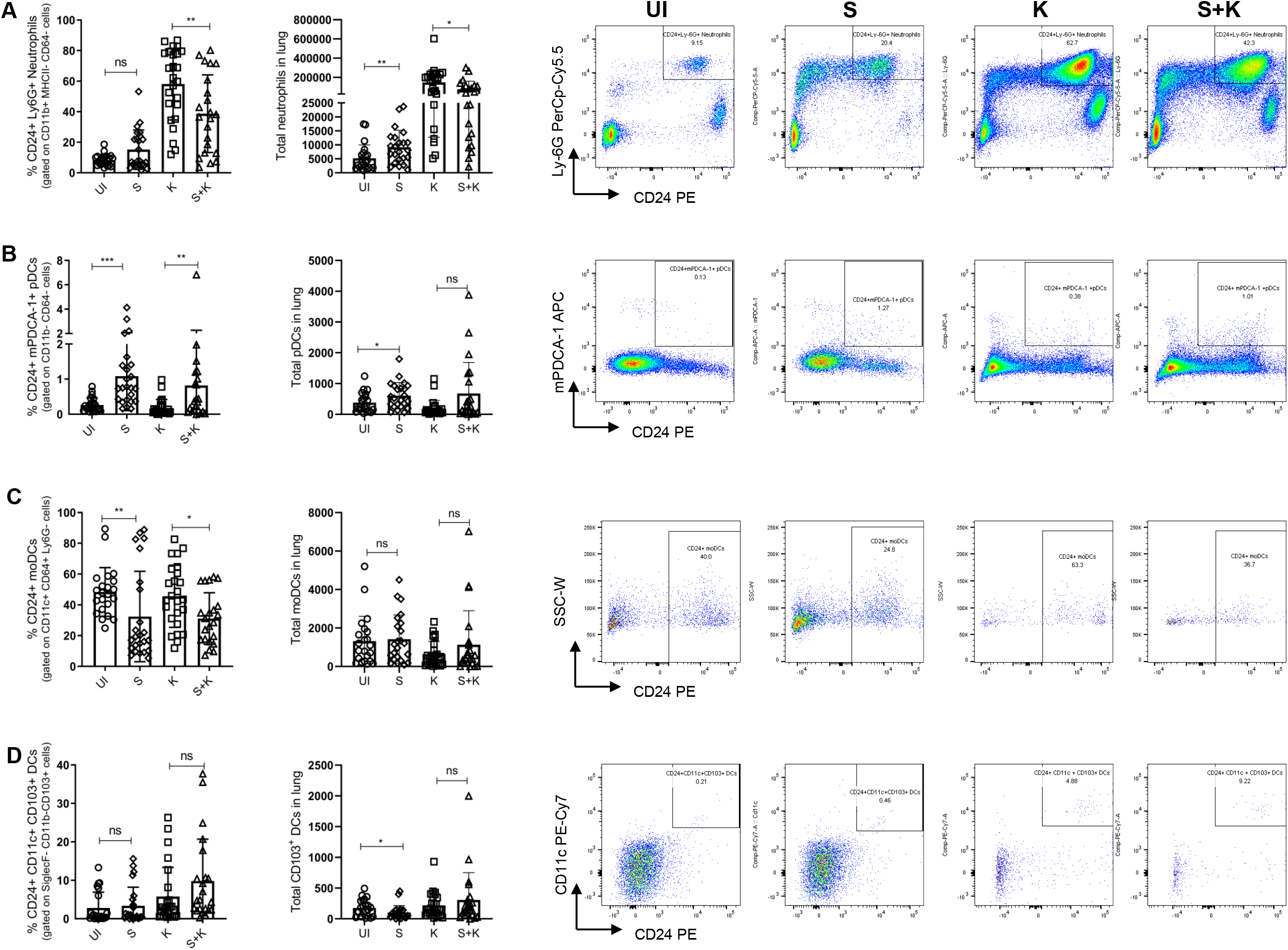
Mice with prior *ST* intestinal infection have altered frequencies of innate cell types in the lung, and altered lung innate cellular responses to respiratory *KP* infection. Percentage frequencies, total numbers and representative FACS plots of (A) neutrophils, (B) lung plasmacytoid dendritic cells (pDCs), (C) monocytic dendritic cells (moDCs), (D) CD103+ DCs are shown in uninfected mice (UI), *ST* infected mice (S), *KP* infected mice (K) and both *ST* and *KP* infected mice (S+K). Data represents cumulative results of five independent experiments (n = 22 for UI, n = 24 for S, n = 25 for K and n = 22 for S+K). Error bars represents mean ± SD and statistical significance difference between UI and S or between K and S+K was determined using Mann-Whitney tests.

We also investigated innate-like T cells including mucosal-associated invariant T (MAIT) cells, invariant natural killer T (iNKT) cells, γδ T cells, and natural killer cells (NK) as they are known to play an important role in bacterial infections (gating strategy in Fig S5B) [10]. No differences were observed in percentage frequencies or total number of iNKT cells, MAITs, γδ T cells or NK cells between mice with and without prior intestinal infection (Fig 5). Furthermore, when complete blood counts were assessed with a Hemavet analyzer, no significant differences were observed in circulating neutrophils, lymphocytes, monocytes, eosinophils, basophils and platelets between mice with and without prior intestinal infection (Fig S7).

**Figure 5:**
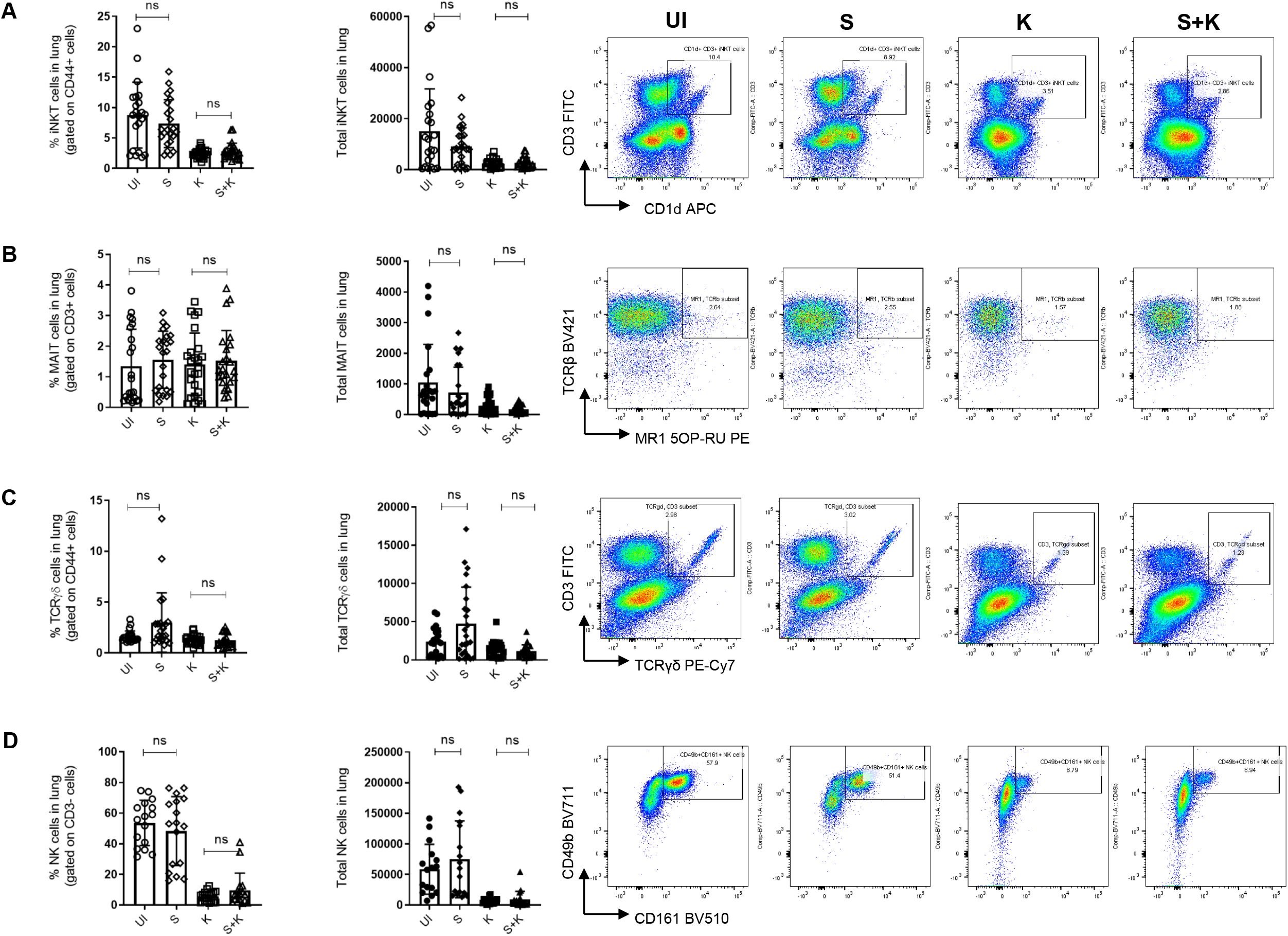
Mice with prior *ST* intestinal infection have no differences in iNKT, MAIT, TCRγδ and NK cells and have increased NET formation in lungs from subsequent respiratory *KP* infection, compared to those with no prior intestinal infection. Percentage frequencies, total numbers and representative FACS plots of (A) iNKT cells, (B) MAIT cells and (C) TCR γδ cells and (D) NK cells are shown in uninfected mice (UI), *ST* infected mice (S), *KP* infected mice (K) and both *ST* and *KP* infected mice (S+K). Data represents cumulative results of five independent experiments (n = 22 for UI, n = 24 for S, n = 25 for K and n = 22 for S+K). Error bars represents mean ± SD and statistically significance differences between UI and S or between K and S+K were determined by using Mann-Whitney tests. (E) Using confocal microscopy, NET formation was assessed in all four groups and images were taken at 60X magnification. Blue fluorescence = nuclear DNA; green fluorescence = citrullinated histone H3; red fluorescence = MPO; NETs are highlighted by yellow arrows in the overlay images. (F) Bar graph showing number of NETs crossing grid lines per high power field. One-way ANOVA with Tukey’s post hoc testing. *** denotes *p* < 0.001.

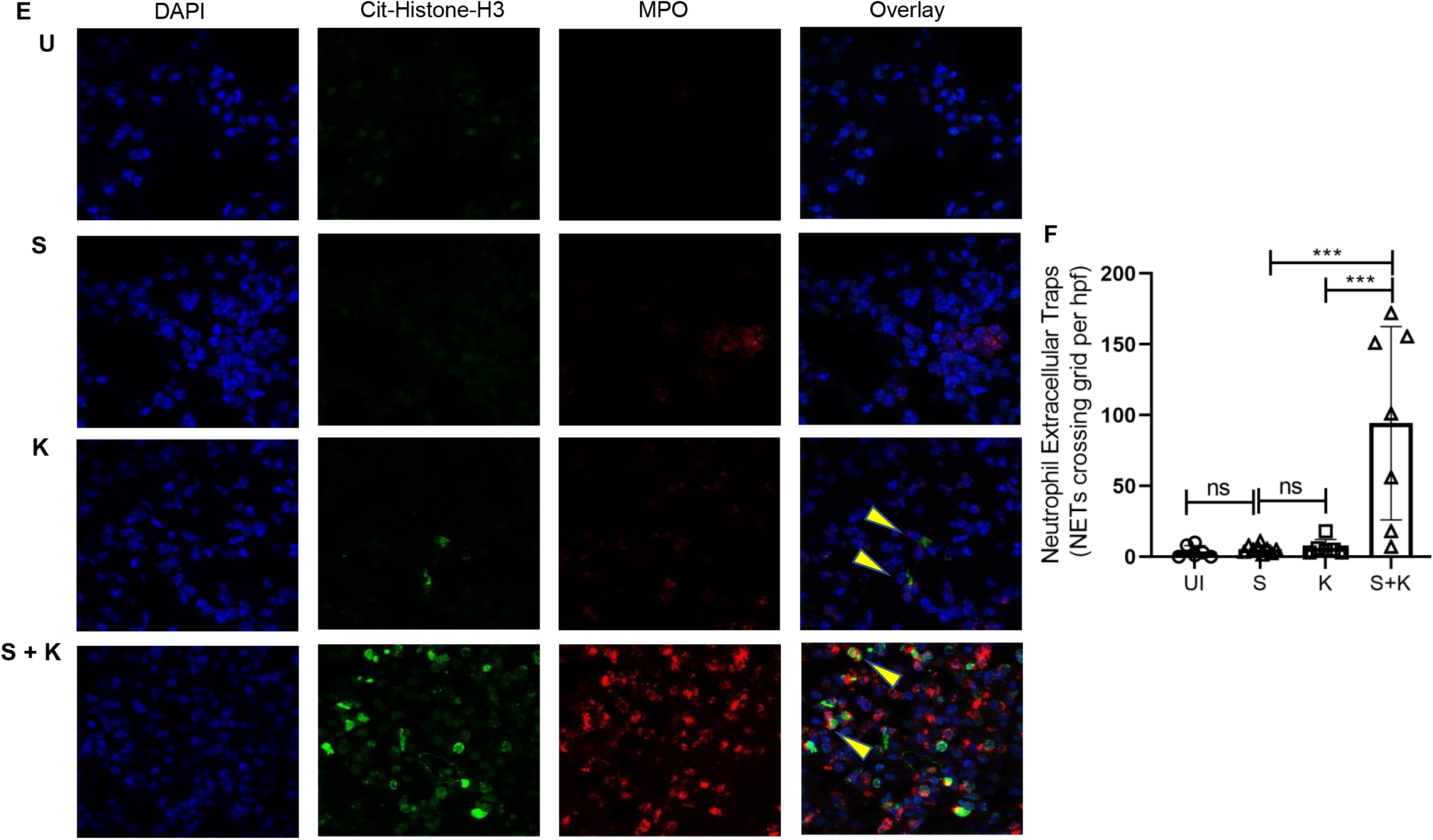

### 5. Mice with prior *ST* intestinal infection demonstrate widespread lung NETosis after *KP* challenge

Recent studies have revealed that excessive NET formation plays a role in pathogen-associated lung injury, including in models of bacterial pneumonia [16–19]. Our histopathological analysis revealed clusters of pyknotic neutrophils and thrombus formation in mice with prior intestinal infection (Fig 2 and Fig S3). It is known that neutrophils constitute a key cellular component of thrombi, and participate in thrombosis by releasing NETs [20]. Moreover, we detected higher levels of IFN-γ in mice with prior intestinal infection compared to mice without prior intestinal infection (Fig 3), and IFN-γ can promote NET formation by neutrophils [21]. We therefore determined whether prior intestinal *ST* infection was associated with increased lung NET formation following intranasal *KP* challenge. We examined citrullinated histones and MPO in lung tissues of each experimental group by immunofluorescence. We found a significantly higher number of NETs (*p* = 0.0006) in mice with prior intestinal *ST* infection compared to those without prior intestinal infection (Fig 5 E and F), as demonstrated by the presence of extracellular DNA overlaid with citrullinated histone H3 and MPO (Fig 5E). Mock-infected lungs did not show any staining for citrullinated histone H3 or MPO. We detected minimal to no NETs in lung tissues of mice in the uninfected control, intestinal *ST* infection alone, and *KP* infected alone groups, and there was no significant difference between these groups (Fig 5F).

## Discussion

We found that bacterial intestinal infection in mice adversely impacts immunity in the lung, increasing susceptibility to secondary respiratory infection. We show that mice with prior intestinal *ST* infection have higher lung bacterial burden and sickness scores after subsequent *KP* challenge compared to mice without prior *ST* infection. This finding was associated with changes in innate cellular responses, most notably those of neutrophils, which were decreased in the parenchyma, clustering in the lung vasculature, and associated with increased NET formation in those with secondary infection. As neutrophils are essential for pulmonary clearance of bacterial infections such as *KP* [22], it is possible that intestinal infection impairs the recruitment and function of lung neutrophil responses against *KP*, leading to an inability to clear the lung bacterial infection, thereby worsening lung function.

In addition to epidemiological studies showing a higher susceptibility to pulmonary infections after intestinal infection in children [2, 3], and that intestinal diseases are often associated with pulmonary disorders [23], the immunological crosstalk of the lung-gut axis is not well understood. In the context of IBD, it has been proposed that intestinal inflammation and increased cytokine levels create conditions favorable for neutrophil margination onto the lung endothelium [24]. When the lung encounters a secondary insult, neutrophil recruitment, activation and extravasation could mediate lung tissue injury and IBD-induced respiratory diseases [8, 25]. Our knowledge of how intestinal infection impacts the recruitment, extravasation and function of neutrophils in tissues outside of the intestine is limited. Studies have shown that GM-CSF plays an important role in neutrophil accumulation [26, 27], and GM-CSF is protective in preclinical models of pneumonia-associated lung injury [28]. We found a significantly lower lung GM-CSF response to *KP* infection in mice with prior intestinal infection, which may decrease the lung’s ability to recruit circulating neutrophils, resulting in an increased susceptibility to infection. In line with this, we also detected increased NET formation and microthrombosis in *KP* infection in mice with prior intestinal *ST* infection as compared to mice without prior intestinal infection. It is likely that NETs and NET-associated factors, including histones and granule proteases, mediate vascular and tissue injury and contribute to microthrombosis [11, 29–31]. The activation of NETosis, the regulated cell death process leading to NET formation, also causes changes in neutrophil morphology including cell membrane rupture and neutrophil death [32]. We speculate that prior intestinal infection induces NET formation contributing to lower numbers of viable neutrophils in the lungs. Further studies examining the mechanisms governing neutrophil activation and NET formation in lungs after intestinal infection are warranted.

Nonconventional T lymphocytes including MAIT cells, *i*NKT cells and γδ-T cells, have tissue-homing properties, and have been implicated in protection against respiratory bacterial infections [7]. Here, we found no differences in frequencies or number of these cells between the groups tested. Likewise, except for IFN-γ we did not find any differences in cytokines that have been implicated in lung defense against bacterial pathogens, including IL-17A [33], TNF-α [34] and IL-10 [35]. Both a protective [36] and detrimental [37] role of IFN-γ has been reported in bacterial infections, and we found significant increases in lung IFN-γ production in mice with intestinal infection. Numerous studies have shown that IFN-γ produced by activated CD4^+^ and CD8^+^ T cells and NK cells plays an important role in protective immunity to *Salmonella* [38, 39]. Although we did not observe differences in NK cell numbers in mice with and without prior intestinal infection, it is possible that activated T cells responding to conserved microbial epitopes are recruited early to the lung after intestinal infection and are a major source of IFN-γ production. However, the importance of T cell activation and IFN-γ production in our co-infection model is unclear and merits further investigation. Modulation of respiratory DCs during *KP* infection has been reported before by Hackstein *et al*. [40]. Although we did not observed differences in total number of DCs, we observed higher frequencies of pDCs in lungs of mice with prior intestinal infection, and this may be related to the role of respiratory pDCs in tissue repair [41]. Alternatively, accumulation of pDCs in lungs post intestinal infection may have involvement in the initiation of inflammation and antigen-specific T cell responses [42]. Moreover, frequencies of moDCs, which may contribute to control of secondary respiratory *KP* infection, was decreased in mice with prior intestinal infection [43]. Further studies are necessary to address the functional role of pDCs and moDCs during bacterial pneumonia after intestinal infection.

Our study has several limitations. Firstly, we did not account for the effect of the pre-existing intestinal microbiota; however, all mice were purchased from the same source, and were cohoused prior to infection. Secondly, we did not evaluate serum levels of IL-6, TNF-α, IFN-γ and VEGF [24] and neutrophil chemokines such as keratinocyte-derived chemokine (KC), macrophage inflammatory protein 2, CXC receptor 2 and CXC ligand 5 [44] which would further our understanding of lung neutrophil trafficking following intestinal inflammation. Thirdly, we have not investigated the role of innate lymphoid cells, which have been shown to be recruited from gut to the lungs in response to inflammation [45]. Fourthly, we have not tested the impact of lower/graded doses of *KP* challenge in mice with prior intestinal infection. Lastly, we emphasize that our study lacks experiments aimed at determining the mechanisms leading to increased lung pathology following intestinal infection. We hypothesize that in secondary infections, neutrophil response and NET production requires a fine balance: underactivity can lead to risk for increased pathogen replication and invasion, whereas over-activity can result in excessive inflammation and tissue damage. Future experiments examining the effect of neutrophil depletion and the effect of NET inhibitors and NET dismantling agents such as DNase are warranted.

In conclusion, the present study demonstrates that infection in the gut adversely impacts immunity in the lung. This report opens up potential avenues for investigating the immunological crosstalk between the lung and gut during enteric infection. While epidemiological studies have demonstrated this lung-gut association, we provide here novel findings that intestinal infection modulates neutrophil and cytokine responses in the lung, resulting in an increased susceptibility to a secondary pneumonia challenge. These data have the potential to inform efforts to prevent and treat respiratory infections in those with intestinal infection or inflammation.

## Acknowledgements

We would like to thank Cole Anderson, Michael Graves, Alexandra Heitkamp and Melanie Prettyman for their technical assistance.

## Notes

### Author contributions

S.T. and D.T.L. designed and directed the project. S.T., O.J. and M.J.C. carried out the experiments. D.T.L., A.H.G., K.J.W., C.C.Y. contributed to data analysis. S.T. and D.T.L. wrote the paper. P.H.S. and C.L. conducted metagenomic next generation sequencing. T.W. analyzed metagenomic sequencing data. All authors discussed the results and commented on the manuscript.

### Financial support

This work was supported in part by grant W81XWH-17-1-0109 from the Department of Defense (to D.T.L.), and by the National Institutes of Health (R01HD093826 to C.C.Y. and R01AI130378 to D.T.L.). This work was supported by the University of Utah Flow Cytometry Facility in addition to the National Cancer Institute through Award Number 5P30CA042014-24, and CMC animal facility.

### Potential conflicts of interest

C.C.Y. authors a US patent (patent no. 10,232,023 B2) held by the University of Utah for the use of NET-inhibitory peptides for the “treatment of and prophylaxis against inflammatory disorders,” for which PEEL Therapeutics, Inc. holds the exclusive license.

## Notes

### Summary of Updates

List of authors updated; Manuscript text and Figures revised and supplemental files updated.

