## Supplementary figures and images for "Intestinal infection results in impaired lung innate immunity to secondary respiratory infection"

### Fig S1

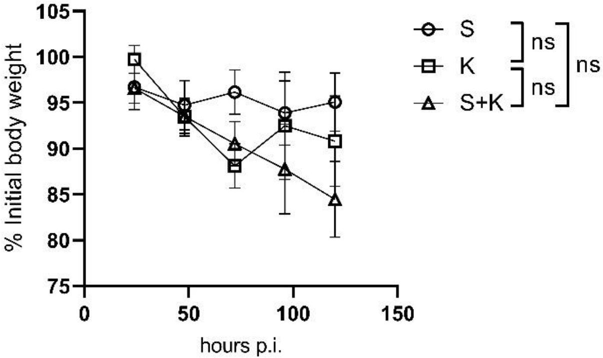

### Fig S2

Uninfected

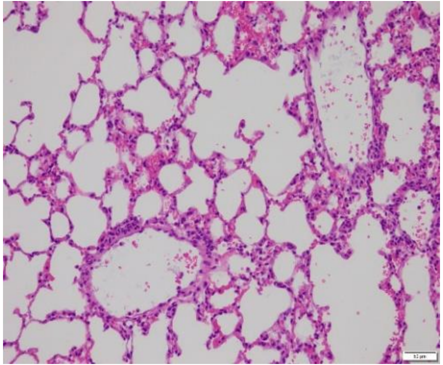

S

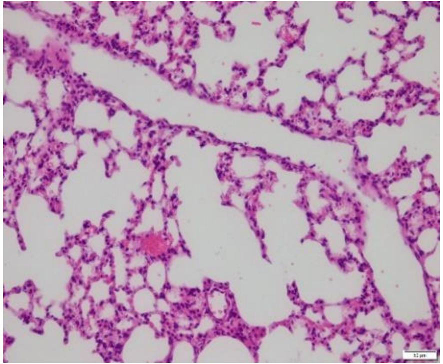

K

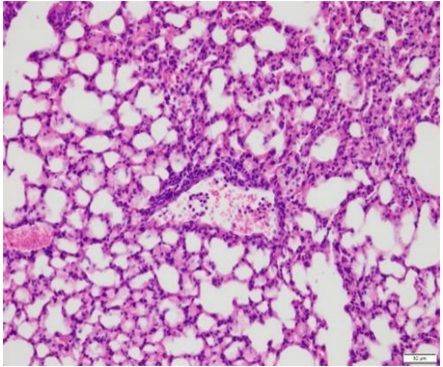

### Fig S3

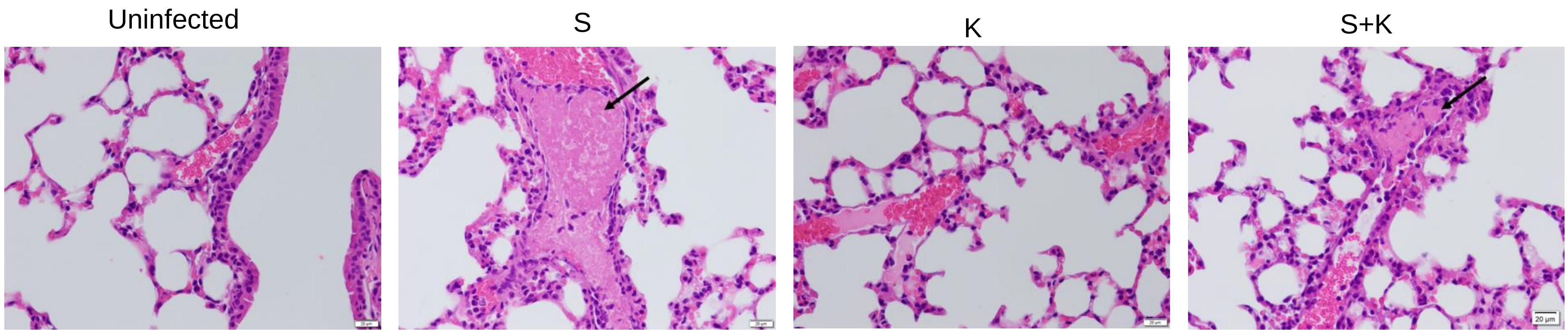

### Fig S4

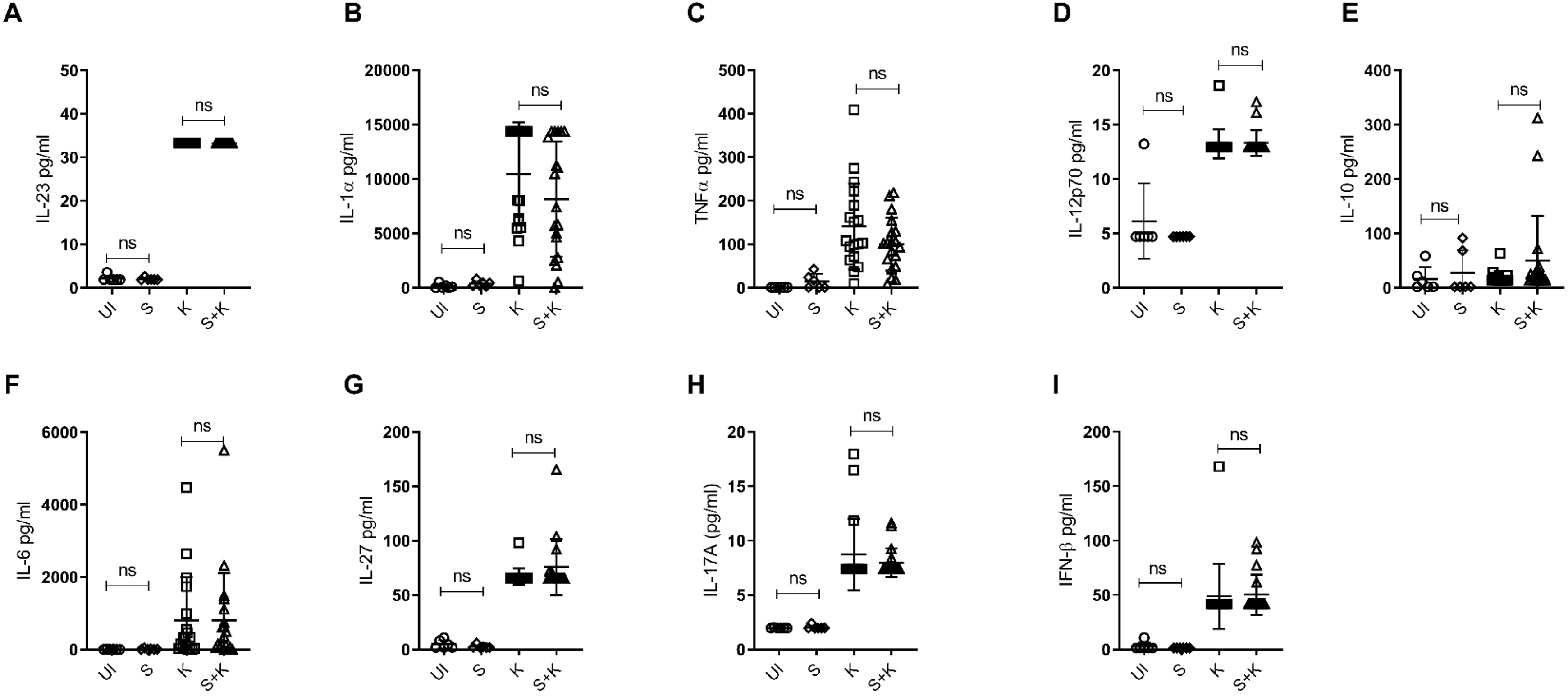

### Fig S5

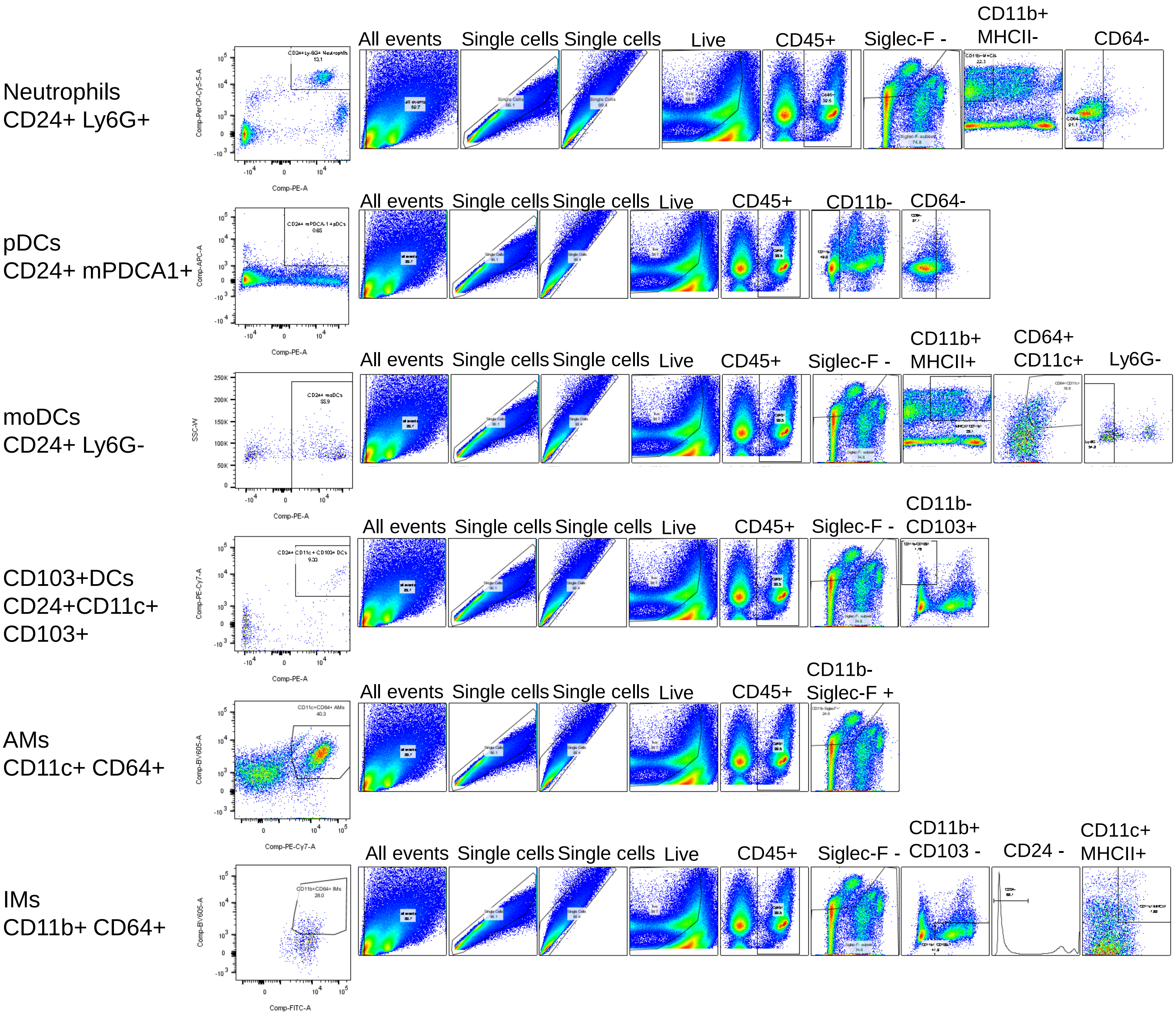

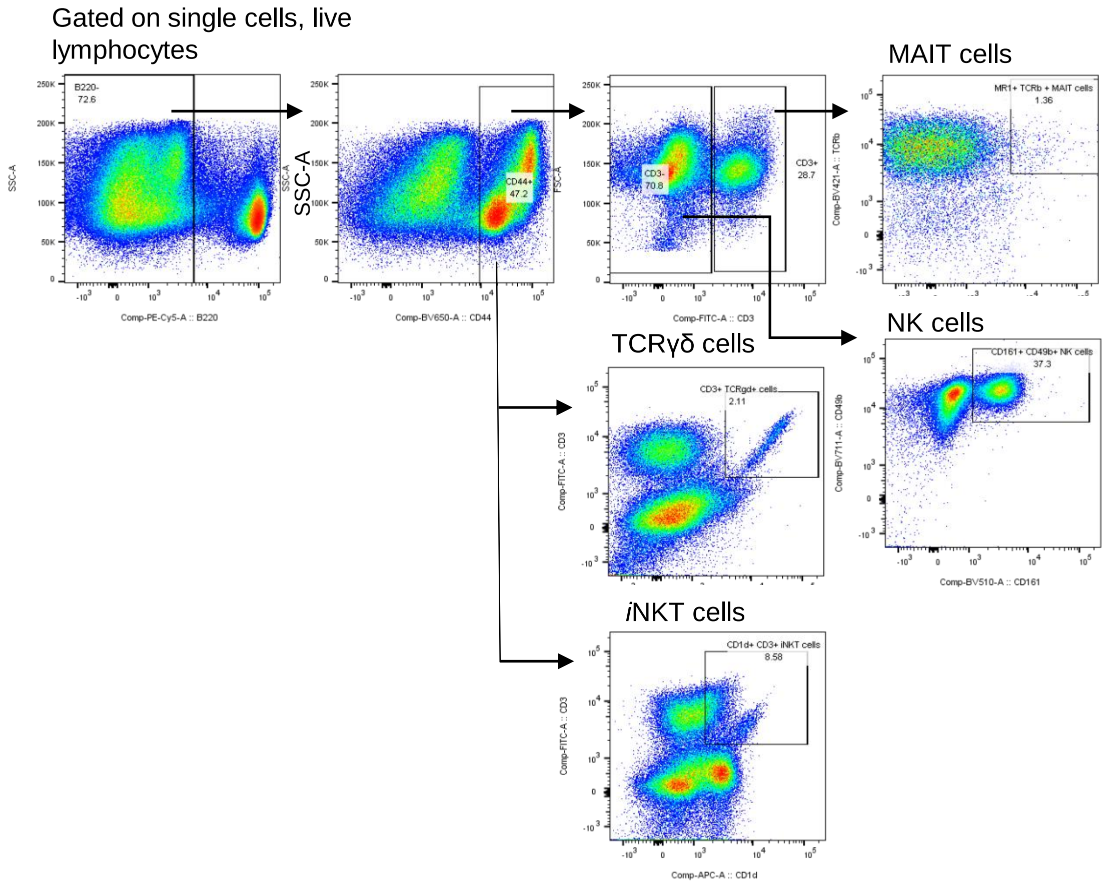

### Fig S6

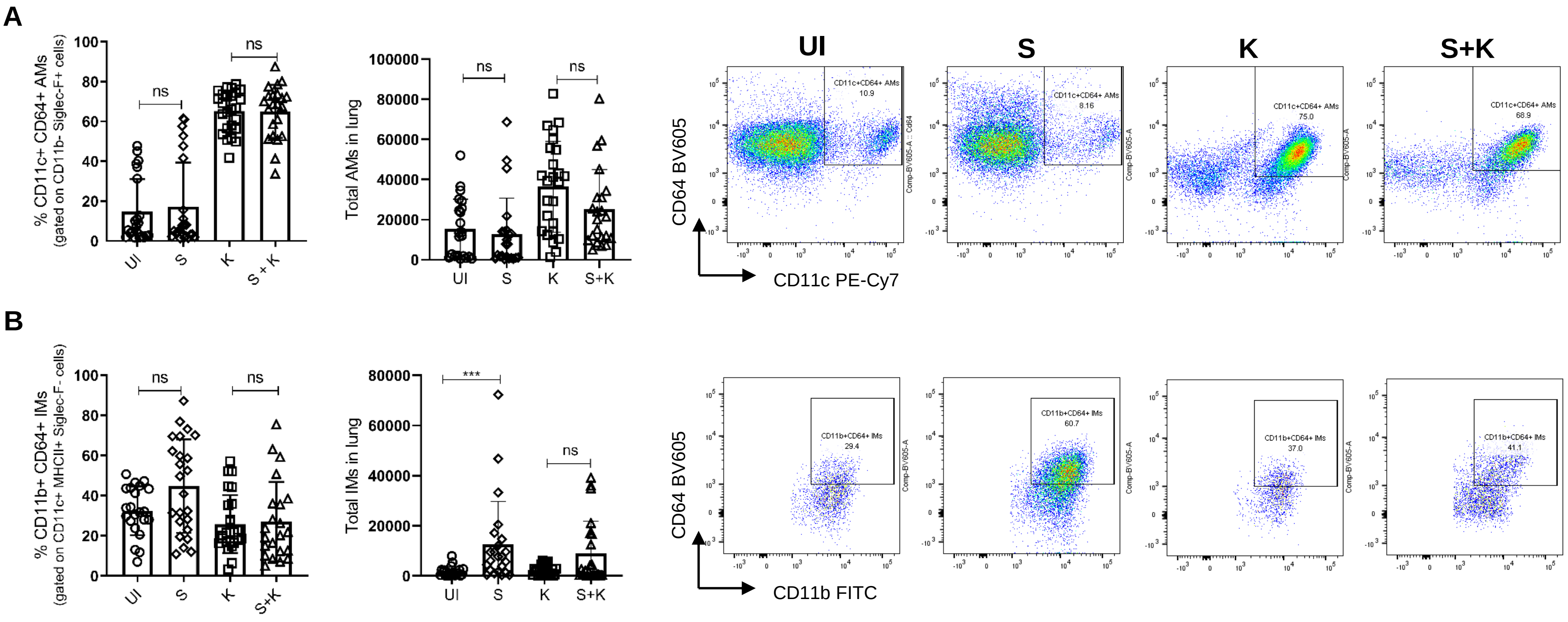

### Fig S7

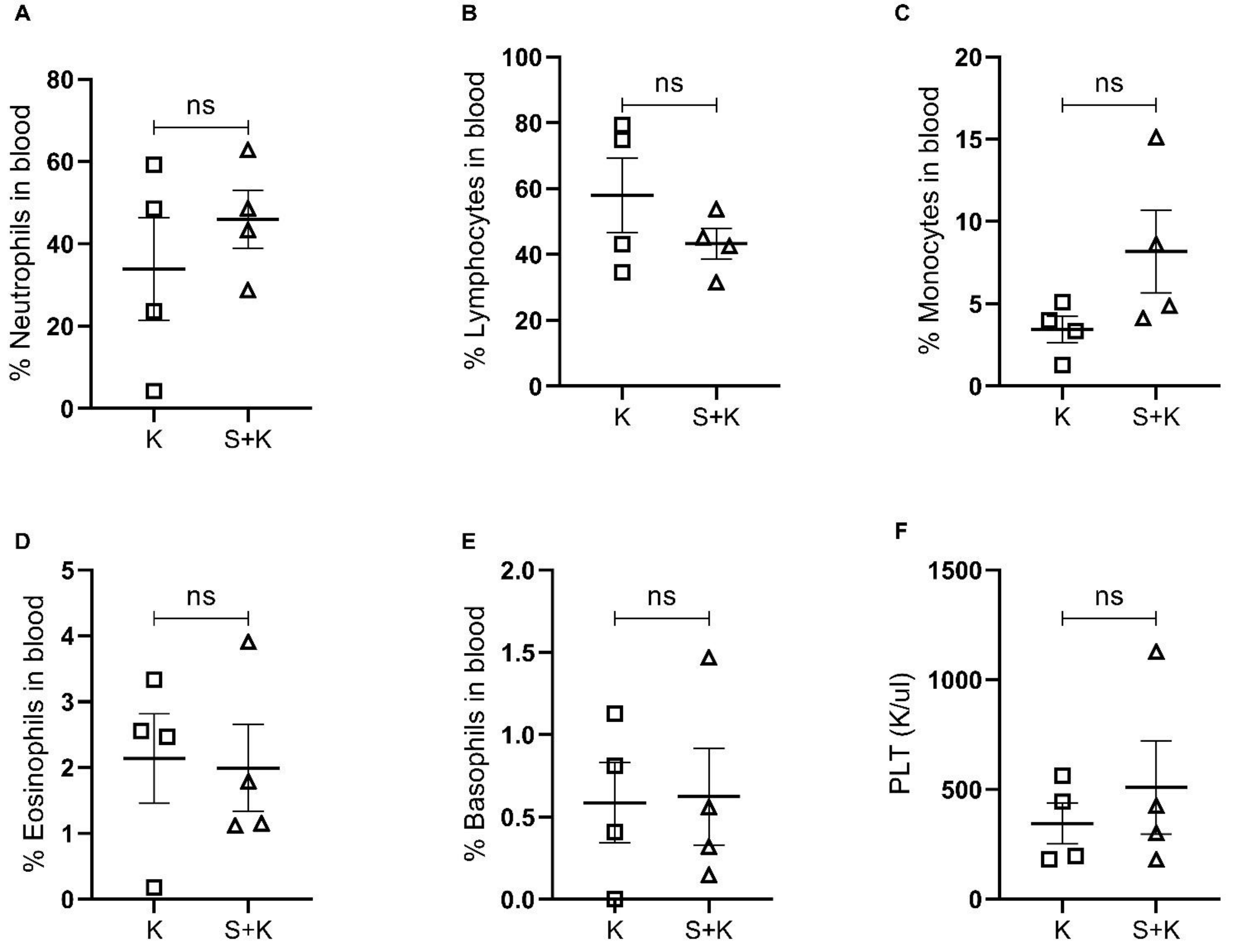
