## Supplementary material for "Intestinal infection results in impaired lung innate immunity to secondary respiratory infection": Table S1

**TABLE S1** Whole Genome Sequencing read alignments from stool and bronchoalveolar lavage (BAL) fluid.

| **Stool Samples** | ***S. enterica* Alignments**[a] |
|---|---|
| S + K[b] 1 | 115,553 |
| S + K 2 | 6,113 |
| S + K 3 | 68,689 |
| S + K 4 | 122,941 |
| S + K 5 | 4,482,383 |
| S + K 6 | 954,438 |
| S + K 7 | 960,685 |
| S + K 8 | 191,777 |
| K[c] 1 | 1 |
| **BAL Fluid** | ***K. pneumoniae* Alignments**[d] |
| S + K 1 | 690,654 |
| S + K 2 | 276,919 |
| S + K 3 | 7,724 |
| S + K 4 | 4,623 |
| S + K 5 | 18,994 |
| S + K 6 | 92,166 |
| S + K 7 | 57,804 |
| S + K 8 | 20,058 |
| S + K 9 | 58,710 |
| S + K 10 | 26,987 |
| K 1 | 88,167 |
| K 2 | 28,446 |
| K 3 | 10,486 |
| K 4 | 17,423 |
| K 5 | 11,538 |
| No DNA Control | 11 |

[a]reads aligned to *S. enterica* LT2 genome using bowtie2
[b]Mice treated with *S. enterica* LT2 and *K. pneumoniae* KPPR1
[c]Mice treated with *K. pneumoniae* KPPR1
[d]reads aligned to *K. pneumoniae* genome using bowtie2
