## Supplementary material for "Intestinal infection results in impaired lung innate immunity to secondary respiratory infection": Table S2

**TABLE S2** Proportion of mice with pyknotic neutrophil clusters (highlighted with arrows in Figure 2A for the S + K group)

| Group | Pyknotic neutrophil clusters |
|---|---|
| Uninfected | Not identified |
| S | Not identified |
| K | Focally present in 2/9 mice, not identified in 7/9 mice |
| S + K | Abundant in 5/9 mice, not identified in 4/9 mice |
