## Supplementary methods for "Intestinal infection results in impaired lung innate immunity to secondary respiratory infection"

**Bacterial strains**

The LT2 strain of *Salmonella enterica* serovar Typhimurium (*ST*; provided by Dr. June Round) was used in all experimental infections. KPPR1 is a rifampicin‐resistant derivative of a *K. pneumoniae* subspecies *pneumoniae*; a clinical pneumonia isolate featuring a type 1 O antigen and a type 2 polysaccharide capsule (ATCC 43816). *ST* was grown overnight at 37°C in Luria Broth (LB) supplemented with 100 µg/ml ampicillin and *KP* was grown overnight in Tryptic Soy Broth at 37°C in a shaking incubator. Overnight cultures were diluted 1:10 in fresh medium and sub-cultured for 4 h under mild aeration. Bacteria were washed twice in phosphate-buffered saline (PBS) and then suspended in 1 ml PBS. *ST* culture (OD_600_ = 0.1) was further diluted 1:10^4^ in PBS and given to mice in a 100 µl volume. *KP* (OD_600_ = 0.4) culture was further diluted 1:10 and given to mice in a 50 µl volume. *KP* bacterial colony forming units (CFUs) were confirmed by culturing on MacConkey agar plates.

Dosing of *S.* Typhimurium was based on titration experiments showing that at the 10^4^ CFU dose, bacteria could be cultured from stool on LB+amp plates but not from homogenized lungs, indicating that *ST* infection did not spread to lungs. Dosing of *K. pneumoniae* for survival studies was based on prior studies in C57BL/6 mice, where varying doses of *KP* via intranasal inoculation yielded a 50% lethal dose (LD50) value of 3 X 10^4^ CFU at 12 days [1, 2] . Based on these studies and to reduce animal pain and respiratory distress, we chose *KP* dose of ~ 10^5^ CFU for survival studies. Dosing of *K. pneumoniae* for study of innate immune response, based on available literature, we chose a high inoculum of 10^10^ CFU to yield an innate immune response within 18 hours [3]

**Tetramer and antibody surface-staining of lung single cell suspensions**

Cells were stained for 30 min at RT with appropriately diluted PE conjugated MR-1-5-OP-RU tetramers or α-GalCer (PBS-44)–loaded CD1d tetramer conjugated to APC, anti-CD3-FITC (Biolegend), anti-CD161-BV510 (Biolegend), anti-CD49b-BV711 (BD), anti-TCRγδ-PE-Cy7 (Biolegend), anti-TCRβ-BV421 (Biolegend), anti-CD45R-PE-Cy5 and anti-CD44-BV650 (Biolegend). To evaluate antigen presenting cells, after Fc Block incubation, cells were also surface stained with anti-CD45 AF700 (Biolegend), anti-CD11b-FITC (Biolegend), anti-CD11c-PE Cy-7 (BD), anti-Siglec-F- BV711 (BD), anti-CD64-BV605 (Biolegend), anti-CD24-PE (Biolegend), anti-Ly-6G-PerCP-Cy5.5 (Biolegend), anti-CD103-BV510 (Biolegend), anti-mPDCA-1 APC (Miltenyi Biotec) and anti-MHC II BV421 (Biolegend) for 30 min at 4 °C

**Metagenomic next-generation sequencing**

Stool and BAL samples were collected in Zymo DNA/RNA shield and DNA was extracted using ZymoBIOMICS™ DNA/RNA Miniprep Kit and quick DNA/RNA kits according to manufacturer’s instructions. DNA yield was determined using the Qubit Fluorometric Quantitation (Invitrogen, USA) at the University of Utah. Sequencing libraries constructed from 100 ng of DNA derived from S + K and K mice underwent paired-end Illumina sequencing according to established methods [4]. Paired-end Illumina reads were aligned with bowtie using default parameters [5] against either the *K. pneumoniae* subsp. pneumoniae strain ATCC 43816 KPPR1 (NCBI accession CP009208.1) genome or the *S. enterica* subsp. enterica serovar Typhimurium str. LT2 (NCBI accession NC_003197.2) genomes.
